# Aquaporin-4 dependent glymphatic solute transport in rodent brain

**DOI:** 10.1101/216499

**Authors:** Humberto Mestre, Benjamin T. Kress, Wenyan Zou, Tinglin Pu, Giridhar Murlidharan, Ruth M. Castellanos Rivera, Matthew J. Simon, Martin M. Pike, Benjamin A Plog, Anna L. R. Xavier, Alexander S. Thrane, Iben Lundgaard, John H. Thomas, Ming Xiao, Aravind Asokan, Jeffrey J. Miff, Maiken Nedergaard

## Abstract

The glymphatic system is a brain-wide metabolite clearance pathway, impairment of which in post-traumatic and ischemic brain or healthy aging is proposed to contribute to intracerebral accumulation of amyloid-β and tau proteins. Glymphatic perivascular influx of cerebrospinal fluid (CSF) depends upon the expression and perivascular localization of the astroglial water channel aquaporin-4 (AQP4). Prompted by a recent publication that failed to find an effect of *Aqp4* knockout on perivascular CSF tracer influx and interstitial fluid (ISF) tracer dispersion, four independent research groups have herein re-examined the importance of *Aqp4* in glymphatic fluid transport. We concur in finding that CSF tracer influx, as well as fluorescently-tagged amyloid-β efflux, are significantly faster in wild-type mice than in three different transgenic lines featuring disruption of the *Aqp4* gene and one line in which AQP4 expression lacks the critical perivascular localization (*Snta1* knockout). These data validate the role of AQP4 in supporting fluid and solute transport and efflux in brain in accordance with the glymphatic system model.

---

A brain-wide fluid transport pathway, known as the glymphatic system, supports the rapid exchange of cerebrospinal fluid (CSF) and interstitial fluid (ISF) along perivascular pathways (*1*). The glymphatic system consists of three principal sequential anatomic segments: (i) CSF inflow along the perivascular spaces surrounding penetrating arteries, (ii) dispersion of CSF through the wider interstitium, and (iii) efflux of ISF along the large-caliber draining veins to re-enter the CSF within the ventricular and cisternal compartments (2). Ultimately, interstitial solutes cleared to the CSF exit the brain through meningeal lymphatic vessels flanking the venous sinuses, along cranial and spinal nerve sheathes, and across the cribriform plate (*3, 4*). Astrocytic endfeet ensheath the cerebral vasculature, and the abundantly expressed astroglial water channel aquaporin-4 (AQP4) localizes primarily to the perivascular endfoot membrane domain abutting the basal lamina. As this anatomic arrangement provides a route for rapid water movement between the perivascular space and the glial syncytium, AQP4 has been proposed to support perivascular fluid and solute movement along the glymphatic system (5).

Several research groups have independently shown that the astrocytic AQP4 is essential for fast glymphatic transport. Illiff et al. (2012) demonstrated a significant suppression of both perivascular CSF tracer influx and interstitial inulin and amyloid-β (Aβ) clearance in *Aqp4* knockout (KO) mice (1). Subsequent work demonstrated that *Aqp4* gene deletion exacerbated glymphatic pathway dysfunction after traumatic brain injury (TBI) and promoted the development of neurofibrillary pathology and neurodegeneration in the post-traumatic brain (6). Plog et al. (2015) similarly found that *Aqp4* KO mice exhibit slowed transport of interstitial solutes to the blood after TBI, reflected by a significant reduction in plasma biomarkers of TBI, including GFAP, neuron specific enolase, and S100β (7). Xu et al. (2015) reported that deletion of *Aqp4* exacerbated Aβ plaque accumulation and cerebral amyloid angiopathy in the APP/PS1 murine model of Alzheimer’s disease (8). Achariyar et al. (2016) found significantly reduced distribution of FITC-ApoE3, ^125^I-apoE2, ^125^I-apoE3, ^125^I-apoE4, as well as ^14^C-inulin in *Aqp4* KO mice following tracer infusion to the CSF (2015) (9). Lundgaard et al., (2016) reported that glymphatic clearance of lactate was reduced in *Aqp4* KO mice (*10*). Finally, Murlidharan et al. (2016) showed that *Aqp4* KO mice exhibit significantly impaired clearance of adeno-associated viruses (AAV) infused into the ventricles, and concluded that glymphatic transport profoundly affects various aspects of AAV gene transfer in the CNS (*11*).

Against this background, the critical role of AQP4 in supporting perivascular CSF-ISF exchange was recently questioned in a report by Smith et al. (2017) (12), in which the authors failed to detect any reduction in CSF tracer influx into the brain parenchyma of *Aqp4* KO mice compared to wild-type (WT) controls. Because a key element of the glymphatic hypothesis is the role of astroglial water transport in supporting perivascular CSF-ISF exchange, we consider it important to re-examine the role of AQP4 in this process, with an aim to resolve the discrepant reports. Using data generated from four independent laboratories, we have undertaken such a re-evaluation of the effects of AQP4 deletion on perivascular glymphatic exchange. Results of this analysis consistently confirm that *Aqp4* deletion impaired perivascular glymphatic flow relative to that in wild-type mice. Our conclusion is strengthened by the use of three independently generated *Aqp4* KO lines, including the line used by Smith et al. (2017) (*12*), as well as the α syntrophin (*Snta1*) KO line, which lacks AQP4 perivascular localization despite normal expression levels (*13*).

## Discussion

This study evaluated the role of the astrocytic water channel AQP4 in the dispersion of fluorescent tracers or contrast agents using four different approaches by four independent groups. Two of the groups (NMU, URMC) injected CSF tracers in cisterna magna and compared the influx of the tracer in coronal sections prepared from *Aqp4* KO and WT mice, whereas the OHSU group injected contrast agent in cisterna magna and then compared CSF transport by DCE-MRI in *Snta1* KO and WT mice. The group at UNC injected fluorescently-tagged Aβ in striatum and quantified it subsequently in coronal sections prepared from *Aqp4* KO and WT mice. Compilation of data from the four independently conducted experiments consistently showed that deletion of *Aqp4 (Aqp4* KO mice) or mis-localization of AQP4 (through *Snta1* gene deletion) potently suppressed influx of CSF tracers or contrast agents, while also attenuating tracer efflux following intraparenchymal injection to striatum. The observations provide strong and concordant support for the glymphatic model in which astroglial water transport via AQP4 supports the perivascular influx of CSF and efflux of ISF (*1*).

Independent replication studies of biological phenomena are essential, both to confirm novel experimental findings, as well as to extend and refine initial observations and interpretations. The original report from the URMC group on the glymphatic system showed that deletion of astrocytic water channels suppressed both the influx of CSF tracers injected into the cisterna magna and the efflux of tracer infused directly in striatum (1). Follow-up of that finding by the NMU group showed that Aβ accumulation and cerebral amyloid angiopathy were significantly increased in a double-hit murine model with *Aqp4* deletion and Aβ over-expression (8). The URMC/OHSU groups subsequently reported that deletion of *Aqp4* aggravated neurofibrillary pathology following TBI (6), whereas the UNC group showed that deletion of *Aqp4* significantly reduced clearance of AAV (11). Thus, these reports conceptually address the same idea, although the replications entailed different approaches and methodologies. The dissenting report from Dr. Verkman’s laboratory belongs to the rarer category of replication studies, which (as their title states) seeks to test directly the glymphatic hypothesis and the AQP4-dependence of solute transport in brain. Since their study was intended to gauge the veracity of the original findings, it should properly be expected to follow precisely the experimental design and data analysis of the original study (*14*), and their methodology should be presented in sufficient detail to allow the reader to assess the fidelity of their replication (*15*). However, Smith et al. (2017) (12) failed to meet these requirements in several key respects. First, their methodologies deviate in critical ways from the original report. In particular, Iliff et al. used ketamine/xylazine anesthesia rather than isoflurane (1). Xylazine is an α_2_ adrenergic agonist that blocks release of norepinephrine from the locus coeruleus projections across the neuroaxis. This is an important point, because norepinephrine has been identified as the key humoral regulator of glymphatic solute transport (*16*), and noradrenaline depletion attenuates glymphatic function. However, with the exception of the photo-bleaching experiments carried out only in WT mice, Smith et al. (*12*) anesthetized their mice with avertin, an injectable anesthetic with an unknown mechanism of action. Avertin is not approved for use in several European countries and is restricted to terminal procedures in the US (*17, 18*). The importance of the anesthetic agent in the present context was highlighted by Groothius et al., (*19*), who reported that parenchymal solute efflux was 100-fold slower rats anesthetized with pentobarbital than those that received ketamine/xylazine, so we suppose that avertin may have had unpredictable effects in this paradigm. Of note, the perivascular influx observed in ketamine/xylazine anesthetized mice are in close resemblance to what is observed for natural sleep (16). Second, we note that Iliff et al. injected CSF tracers using a 30 GA needle with an inner diameter of 160 μm: their needle was connected to a constant-pressure syringe pump and remained in place for the duration of the experiment (1). This procedure raises intracranial pressure transiently by ~ 2 mm Hg, although a side-by-side comparison showed that this small pressure pulse had no effect on tracer distribution (20). In contrast, Smith et al. used a beveled-glass micropipette with an inner diameter of 20 μm connected to a pressure injector (10 psi, 50-100 ms pulse duration), and did not monitor ICP, raising the possibility that they inadvertently perturbed glymphatic function (12). Third, Iliff et al., based their analysis on mice aged 2-3 months, whereas the age range was 3-6 months in the Smith study (*12*). The wider age range will likely have affected their data, since murine glymphatic influx and clearance both declines steeply with increasing age (21). However, the age distribution within the *Aqp4* KO and WT groups were not reported in the Smith et al., publication. Finally, key methodological details, such as the injection technique and volume, anesthesia, or the age of the animals were not provided for data collected in rats. Notably, the efficacy by which *Aqp4* was deleted in the rat brain was not confirmed in a previous report (*12*).

Since the initial description of glymphatic exchange of CSF and ISF along perivascular spaces (1), several additional related studies have been published (22–24). In light of this increased understanding of fluid movement in the brain, it is critical that we assess how these data fit with the glymphatic hypothesis as it was initially proposed and thus identify elements of the hypothesis that require further investigation. The initial study articulating the glymphatic hypothesis confirmed seminal work by Cserr and colleagues that interstitial tracers are cleared from brain tissue at rates that are independent of molecular weight (*1, 25*). While these early studies support the role of bulk flow in the clearance of interstitial solutes from the brain interstitium, they lacked the spatial resolution necessary to resolve whether bulk flow was occurring throughout the entire brain interstitium, or was restricted to a subset of compartments permissive to flow. A large body of indirect experimental evidence supports the occurrence of bulk flow in the periarterial space in the brain (see the review by Hladky and Bartrand 2014, (*26*) and Nicholson and Hrabetova (*27*)), and there is indirect evidence that this flow is peristaltic, propelled by pulsations of the arterial walls driven by the heartbeat (e.g., Hadaczek 2006, (*28*)). Smith et al. (12) themselves provide evidence for bulk flow in the perivascular space, stating “occasionally, when the injected dextran entered the perivascular space of vessels near the injection site, both large and small dextrans traveled substantial distances away from the injection site” (12). Several fluid-dynamic models and experimental studies have pointed to the existence of convective flow in the periarterial space driven by arterial pulsations (*29–33*). The models generally predict that, although there are local and transient negative velocities (i.e. reversed flow) at times, the time-averaged (bulk) flow is in the direction of the pulse wave, i.e., in the same direction as the arterial blood flow. These studies are supported by in vivo dynamic imaging studies, including MRI experiments in human subjects, demonstrating the rapid distribution of CSF tracers into and through the brain interstitium along peri-arterial routes (*24, 34, 35*). In addition to perivascular spaces, white matter tracts also serve as permissive routes for interstitial solute and fluid movement (36, 37).

The claim by Smith et al. (12) that solute transport in the parenchyma is primarily driven by diffusion rather than convection (bulk flow) is based on their finding of a dependence of this transport on the size of the solute particles. They claim that diffusive transport depends strongly on particle size, whereas convective transport does not, so long as the ratio of the particle size to the pore or channel size is less than about 0.5. In support of this claim, they cite a single reference (Dechadilok and Deen, 2009, (38)), which does not in fact support their claim, but rather shows a significant dependence of flow resistance on particle size for much smaller particles. The influence of solutes or suspended particles on flow through narrow channels is a complicated problem in hydrodynamics, entailing many factors, including solute concentration, electrical charge, shape of the particle, channel geometry, tortuosity, and possible obstructions (*27*). Hence, it shall require development of a very thorough and sophisticated set of theoretical models to exclude bulk flow as a significant contributor to the transport observed experimentally by Smith et al (*12*). However, their findings are in general agreement with several recent modeling studies suggesting that at the microscopic scale encompassing small numbers of vessels and the interposed neuropil, diffusion and dispersion are sufficient to account for experimentally-observed fluid and solute movement (*22, 39, 40*). Notwithstanding, it is important to note that these findings do not exclude a contribution of bulk flow within the entire interstitium, and do not define whether the experimental approaches used to detect flow on the microscopic scale are sufficiently sensitive to detect bulk flow occurring at very low flow rates over long distances. If bulk flow within the interstitium occurs along peri-arterial spaces and white matter tracts, then there must be a sink for this directional flow; conservation of mass (as expressed by the continuity equation) demands that this volumetric flow must be continuous throughout the other parts of the glymphatic system – i.e., in the neuropil and the perivenous spaces. Although the full volumetric net flow must pass through the neuropil, flow speeds will be lower because of the much larger total cross-sectional area of the available extracellular flow channels, and due to the availability of the gap junction-coupled astroglial syncytium as an intracellular route for water and solute movement (41).

As with any scientific hypothesis, the initial description of the glymphatic system articulated by Iliff et al. (1) must undergo refinement as new experimental data or insights from modeling experiments become available. As discussed above, animal experiments, and new studies carried out in human subjects, have consistently supported the importance of perivascular pathways and white matter tracts as permissive routes for bulk flow through brain. These pathways, which connect different brain regions with one another, and link every brain region with the CSF compartments, appear to be the anatomical scaffold that organizes glymphatic fluid and solute exchange throughout the brain. The necessary role for astroglial water transport (e.g. AQP4) in supporting perivascular CSF-ISF exchange, the key innovation that distinguished the glymphatic hypothesis from established thinking about perivascular bulk flow (42), has been supported in the present study by findings from four independent research groups. What remains unclear is whether slow bulk flow takes place within the interstitium outside of perivascular spaces and white matter tracts, or whether local diffusion supported by glial function spans the microscopic gaps between these permissive bulk flow pathways. In addition, the role that other elements of the neurovascular unit, including astrocytes, or other cell types such as pericytes and smooth muscle cells play in perivascular glymphatic exchange remain to be experimentally defined.

Thus, as articulated by Holter et al. in their recent modeling study (*40*), there is a need to develop more appropriate technical approaches for evaluating slow interstitial bulk flow over long distances (*43*), such as DCE-MRI in nonhuman primates or human subjects. Until then, the glymphatic model of CNS fluid and solute flow should properly include the rapid exchange of CSF with the brain ISF along perivascular spaces, with CSF principally entering along peri-arterial spaces and ISF draining towards the ventricular and CSF compartments along peri-venous pathways, linked to one another by AQP4-dependent astroglial water transport. On this last element, the present findings from four independent groups do not concur with the negative findings reported by Smith et al. (*12*). Instead, our compiled data provides strong and consistent support for our claim that solute transport in the rodent brain is facilitated by the polarized expression of AQP4 in astrocytic endfeet.

## Materials & Methods

### Generation of four transgenic mice lines with deficient expression of AQP4 in astrocytic vascular processes

#### NMU lab

*Aqp4* KO mice were generated by Dr. Fan Yan by targeted gene disruption as described previously (44). In brief, an AQP4 replacement targeting vector was constructed using the positive-negative selection cassettes derived from the vectors pPolII long neo bpA and pXhoMC1TK, containing the neoR and HSVtk genes, respectively. The targeting construct was linearized with NotI and introduced into E14K ES cells by electroporation. ES cell clones that were G418/gancyclovir-resistant were isolated, amplified, and screened for targeting fidelity using Southern blot analysis. Cells from two targeted clones were microinjected into CD1 blastocysts and implanted into pseudopregnant recipients (44). Five-month old *Aqp4* KO and WT mice were used in the present study.

#### UNC lab

The constitutive *Aqp4* KO mouse model was generated by Dr. Alan Verkman (*45*). In brief, *Aqp4* KO mice were generated on a CD1 background, following the construction of a targeting vector for homologous recombination of a 7-kb SacI AQP4 genomic fragment, in which part of the exon 1 coding sequence was deleted. All animal experiments reported in this study were conducted on *Aqp4* KO mice (B6/129) or B6/129 controls.

#### URMC

*Aqp4* KO mice were generated by the Dr. Ole Petter Ottersen group using GenOway technique, as described previously (46). Their strategy involved cloning and sequencing of a targeted region of the murine *Aqp4* gene in a 129/Sv genetic background. Identification of a targeted locus of the *Aqp4* gene permitted the delete exons 1–3 to avoid any expression of putative splice variants. Hence, a flippase recognition target (FRT)-neomycin-FRT-LoxP-validated cassette was inserted downstream of exon 3, and a LoxP site was inserted upstream of exon 1 (46). The mice were backcrossed for 20+ generations with C57BL/6N mice prior to experimentation.

#### OHSU

The *Snta1* KO mouse line was generated by Dr. Stanley Froehner (47), and was purchased through The Jackson Laboratory (*Snta1^tm1Scf^/J, #012940*). In brief, exon 1 of the mouse *Snta1* gene was replaced with a neomycin resistance cassette. The resulting construct was electroporated into 129P2/OlaHsd-derived E14 embryonic, and these cells were then injected into C57BL/6J blastocysts. The resulting mice were bred with C57BL/6J mice for at least 11 generations before the colony was established. Adult male and female mice (WT 15.4 ± 1.6 weeks, *Snta1* KO 14.7 ± 1.6 weeks) were used in these studies.

### Glymphatic analysis

#### NMU

##### Intracisternal Tracer Infusions

Intracisternal injections of Texas Red conjugated dextran (Dex-3; 3 kDa, Invitrogen) were adapted from a previous report (1). In brief, the mice were anesthetized by a 4% chloral hydrate solution. A ten microliter volume of 0.5% Dex-3 dissolved in artificial CSF was infused into the cisterna magna through a 50 μL syringe mounted with a 27-gauge needle (Hamilton, Reno, NV, USA), connected to a constant current syringe pump (TJ-2A/L07-2A, Suzhou Wen Hao Chip Technology Co. Ltd., Jiangsu, China). The intracisternal infusion was carried out at a rate of 2 μL/min. Thirty min after the start of infusion, the still deeply anesthetized animals were perfused trans-cardially with 4% paraformaldehyde (PFA), and the brains removed and post-fixed in the same fixative for 24 hours. All experiments were approved by IACUC (Institutional Animal Care and Use Committee) of Nanjing Medical University. All efforts were made to minimize animal suffering and to reduce the number of animals used for the experiments.

##### Tissue processing and imaging analysis of fluorescent tracer

The telencephalon was cut into 100 μm coronal sections using a vibratome (Leica, Wetzlar, Germany). Sections were mounted onto gelatin-coated slides in sequence, then washed and cover-slipped with buffered PBS/glycerol. Images of whole brain section were captured by a digital microscope (Leica Microsystems, Wetzlar, Germany) under a 1.25x objective, with uniform exposure time, offset, and gain. The penetration of Dex-3 into the brain parenchyma was quantified using NIH Image J software as described previously (1). In brief, the fluorescent tracer coverage within the whole section was measured by the interest grayscale threshold analysis with constant settings for minimum and maximum intensities. In addition, tracer penetration along perivascular spaces and into brain parenchyma was quantified on high magnification images of the hypothalamus, which were captured with 10x objective. The fluorescence intensity within the perivascular space were measured along large vessels (diameter >20 μm) extending up to 500 μm below the brain surface. Two or three separate vessels from each image were analyzed and the data averaged. The fluorescence intensity in the brain parenchyma was also determined along corresponding linear region adjacent to the vessels (Fig. 2B). The results were averaged from 6-8 brain slices per mouse and five mice for each genotype group. The imaging capture and analysis was performed by an investigator who was blind to animal genotype.

#### UNC

##### Intracranial Amyloid-β injections

The experiments were performed as described previously (*11*). In brief, postnatal day 0 (P0) mouse pups were anesthetized by placement on ice for two minutes, and then received intrastriatal (ISTR) injections using a stereotaxic apparatus. A one μl portion of hilyte-555 labeled amyloid-β peptide (AS-60480-01 Anaspec) (reconstituted to 1 mg/ml concentration in 1XPBS) was injected into the left striatum (coordinates: 1.25 mm relative to the sagittal sinus, 2 mm rostral to transverse sinus and 1.5mm deep) with a Hamilton 700 series syringe bearing a 33 gauge needle (Sigma-Aldrich, St. Louis, MO), and attached to a KOPF-900 small animal stereotaxic instrument (KOPF instruments, Tujunga, CA). Mice were revived under a heat lamp and rubbed in the bedding (to avoid olfactory cues) after injections before being placed back with the dam. Three hours post injection, mouse pups were rapidly anesthetized using hypothermic shock as above, and perfused with 1XPBS followed by 4% PFA. All experiments were performed in accordance with the guidelines set forth by the University of North Carolina Committee on Animal Welfare.

##### Tissue processing and confocal microscopy

The mouse brains were harvested and post-fixed for 24 hours prior to sectioning. In brief, 50 μm thick coronal plane vibratome sections were obtained using a Leica VT 1200S apparatus (Leica Biosystems, IL). Free-floating mouse brain sections were mounted with ProLong™ Gold Antifade Mounting Media with DAPI (Life Technologies, CA). We used the Zeiss CLSM 700 confocal laser scanning microscope for imaging mouse brain sections (Microscopy services laboratory, UNC). The images were stitched, pseudocolored and analyzed on the Zen^®^ Black software.

#### URMC

##### Intracisternal Tracer Infusions

Mice were anesthetized with ketamine-xylazine (100-10mg/kg) and fixed in a stereotaxic frame whereupon the posterior atlanto occipital membrane overlying the cisterna magna was surgically exposed. For all CSF tracer experiments, Alexa 647-conjugated bovine serum albumin (BSA-647; 66 kDa; 0.5%) was infused into the subarachnoid CSF via cisterna magna puncture, at a rate of 1 μL/min for a period of five min (5 μL total volume) through a 30-gauge syringe pump (Harvard Apparatus). To visualize penetration of fluorescent CSF tracers into the brain parenchyma ex vivo, anesthetized animals were decapitated at 30 min after the start of infusion, a time point previously identified to correspond to robust tracer penetration of similar molecular weight compounds in young male C57BL/6 mice (1). Brains were then removed and post-fixed in 4% PFA for 24 hours before being sliced with a vibrotome into 100 μm coronal sections which were slide-mounted using PROLONG anti-fade gold with DAPI (Invitrogen). All experiments were approved by the University Committee on Animal Resources of the University of Rochester.

##### Ex-vivo imaging of fluorescent CSF-tracers

BSA-647 circulation along perivascular pathways and penetration into the brain parenchyma was visualized by conventional fluorescence microscopy of whole brains and 100 μm vibratome coronal brain sections, as described previously (1). Whole brain images were acquired on a macroscope (MVX10, Olympus) at 12.5x magnification. For coronal sections, six brain sections (per animal were imaged and analyzed by a blinded investigator using an Olympus fluorescence macroscopy (BX51) under 4x objective power to generate whole-slice montages (using the CellSens software, Olympus). Tracer penetration was quantified using NIH Image J software as described previously (1). The first slice was collected at the beginning at the anterior aspect of the corpus callosum, one section was collected every 500 μm apart until a total of six sections had been collected for each animal. Fluorescence of the CSF tracer BSA-647 was measured in each slice as mean pixel intensity (MPI). The MPI from the six brain slices from each animal were averaged to define CSF penetration within a single biological replicate. A subset of brain slices were immunostained for AQP4 as previously described (*21*).

#### OHSU

##### Dynamic contrast enhanced magnetic resonance imaging (DCE-MRI) of glymphatic transport

3-6 month old mice were anesthetized with isoflurane, with induction at 3-5%. The posterior atlanto-occipital membrane was exposed surgically. To enable the contrast infusion during MRI scanning, we used a pulled glass micropipette with trimmed end (external diameter of approximately .016 mm) to perform the infusion. The micropipette was fixed in place for the duration of the imaging session with superglue. Gadoteridol (68 mM, osmotically adjusted), a contrast medium was infused by syringe pump (Harvard Apparatus) at a rate of 500 nl/min for 20 minutes (10 μl total volume), with a 2 μl chase of saline.

##### MR imaging

CSF circulation was quantified by dynamic contrast-enhanced magnetic resonance imaging (DCE-MRI). All imaging was performed using a Bruker-Biospec 11.75 T preclinical scanner equipped with a 20 mm I.D. quadrature RF volume-coil with a specially designed head holder. Heart rate, oxygen saturation and respiratory rate were monitored, and core temperature was maintained at 37 °C using a warm air temperature control system (SA Instruments). Upon placement of the glass micropipette, isoflurane anesthesia was switched to ketamine-xylazine (100-10mg/kg) for the duration of the experiment. Consecutive T_1_ weighted 3D FLASH images were obtained at 10 minute intervals (TR/TE 16/2.8 msec, flip angle 15°, matrix 256×192x192, 100×100x100 μm resolution), for a total of 90 minutes. Infusion was initiated after acquiring the first image. If no elevation in signal was detectable in the cortical or subcortical brain regions or the basal cistern within the first 30 minutes, the imaging session was aborted and the animal was excluded from further analysis.

##### MR data analysis

Due to frequent occurrence of motion artifact during the last 30 minutes of acquisition (70-90 minutes), these three time points were excluded from analysis, and those animals showing movement artifacts during the first 60 minutes of acquisition were also excluded. Linear rigid body registration was used to spatially align the image series to the baseline image (FSL). Regions of interest (ROIs) were drawn on a coronal image slice best matching that depicted in Figure 6I, and mean intensity values were determined with FIJI software. Repeated measures two-way ANOVA with Sidak multiple comparisons correction was used to analyze the data. Multiple comparisons were used to determine the significance of genotype differences at each time point.

### The data that support the findings of this study are available from the corresponding author(s) upon reasonable request

#### Results

The study included data from four laboratories in three countries using three independently generated *Aqp4* KO lines and one *Snta1* KO mouse line (Fig. 1a-b). Immunohistochemistry done in parallel with the glymphatic experiments verified that AQP4 was indeed deleted in all the *Aqp4* KO mouse lines (Fig. 1c). In the *Snta1* KO mice, immunofluorescence demonstrated that perivascular AQP4 localization was absent in this line (Fig. 1c).

**Fig. 1.**
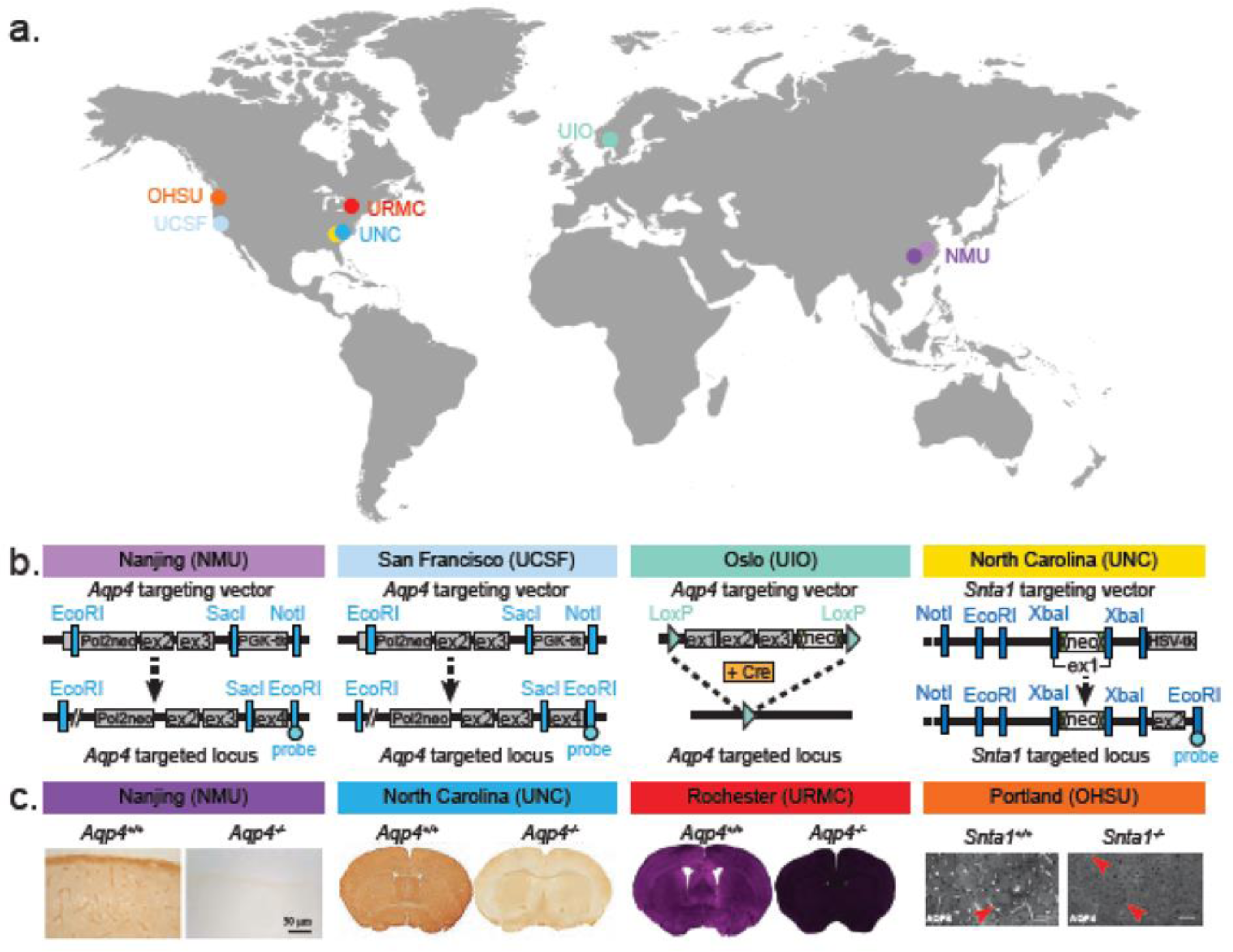
Strategy used for generation of *Aqp4* and *Snta1* KO mice. (a) World map depicting the location of the three labs that generated the *Aqp4* KO mice and the lab generating the *Snta1* KO mice, as well as the four labs that collected data on glymphatic functions in the four transgenic mice lines. Nanjing Medical University (NMU), University of California San Francisco (UCSF), University of North Carolina (UNC), Oslo University (OU), University of Rochester Medical Centerl (URMC), Oregon Health & Science University (OHSU). (b) Diagram depicting the strategy for deleting *Aqp4* or *Snta1.* (c) Immunohistochemical analysis comparing AQP4 expression in WT and *Aqp4* KO mice, as well as immunolabeling showing lack of perivascular AQP4 localization in *Snta1* KO compared to WT mice. The immunohistochemistry was performed in parallel with the functional analysis presented in Figs. 2-5.

### NMU: Reduced glymphatic influx of intracisternal fluorescent tracer, TR-d3, in the Aqp4 KO mouse brain

We injected intracisternally the fluorescent tracer Texas Red-dextran (3 kD, TRd3) and compared overall penetration of TR-d3 in coronal sections in WT and *Aqp4* KO mice, as described previously (*1*). The analysis showed that the fluorescence coverage area was significantly reduced in all five *Aqp4* KO mice compared to WT controls (Fig. 2a-b). A quantitative analysis further revealed that the percentage area with accumulation of TRd3 in *Aqp4* KO brain was only one third of that in WT control (Fig. 2c). High magnification imaging of the hypothalamus revealed that AQP4 KO impaired the influx of TR-d3 into both the perivascular space and the brain parenchyma (Fig. 2b). Quantification of the intensity of TR-d3 as a function of the distance from the brain surface showed rapid decay of tracer with increasing distance from the brain surface in *Aqp4* KO mice: The tracer was almost undetectable at 500 μm below the brain surface in both the perivascular space and in the brain parenchyma in *Aqp4* KO mice. By contrast, the TR-d3 fluorescence intensity remained at 90% of the pial surface intensity to a depth of 500μm in the perivascular space as well as in the adjacent parenchyma in WT mice (fig. 2d-e). Taken together, these results replicated the previous finding that AQP4 facilitates the transfer of intracisternally injected TR-d3 from brain CSF to the parenchyma (1).

**Fig. 2:**
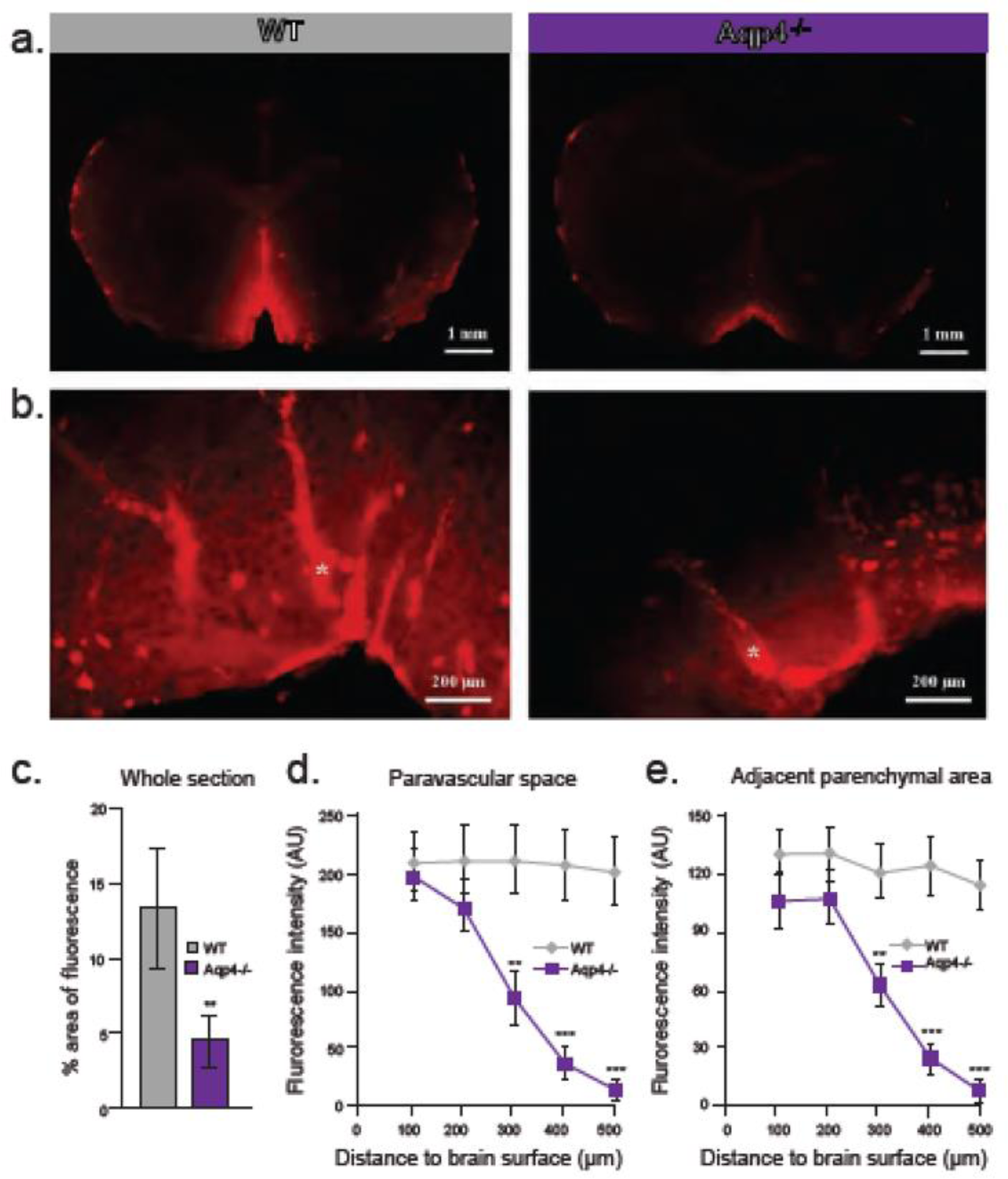
NMU: Aqp4 gene deletion reduced the penetration of intracisternally injected tracer into the brain parenchyma. Texas Red-conjugated dextran (Dex-3, 3kD) was injected intracisternally into WT and Aqp4 KO mice. Thirty minutes after injection, the anesthetized animals were perfusion fixed, and the fluorescence was evaluated ex vivo. (a) Representative images showed that compared to WT mouse brain, CSF tracer penetration into the brains of *Aqp4* KO mice was markedly reduced. (b) Quantification of the percentage area of whole-slice fluorescence of the both genotypes. (c) Representative high magnification micrographs of the hippocampus showed that the fluorescence intensity of Dex-3 within the perivascular space (star) and adjacent brain parenchyma was significantly reduced in *Aqp4* KO mice, relative to WT controls when plotted as a function of increased distance from the brain surface. (d-e) Quantification of Dex-3 penetration showed that the mean fluorescence intensity along the perivascular space and the interstitium adjacent to the vessels was significantly reduced at 300 μm below the brain surface of *Aqp4* KO mice. By contrast, in WT mice, the perivascular and interstitial fluorescence signal remained high to at least 500 μm from the surface. AU, arbitrary units.** *p* < 0.01, student-t test; *Aqp4* KO vs WT, mean ± standard deviation, n = 5 per group

### UNC: Reduced spread of fluorophore-tagged Amyloid-Beta (Aβ) peptide in the Aqp4 KO mouse brain

To probe the effects of AQP4-mediated CSF flux on spreading of pathological CNS deposits, we compared the spread of tracer after intraparenchymal administrations of fluorophore-tagged Aβ peptide within WT and *Aqp4* KO neonatal mice. Specifically, we injected postnatal day 0 (P0) mice with 1 μl hilyte-555 labeled amyloid-ß (Aß) peptide (AS-60480-01 Anaspec). At three hours post-injection, we observed visibly stronger Aß peptide mediated fluorescence signal (magenta) lying close to the striatal site of administration in *Aqp4* KO cohorts, as compared to that seen in WT mice (Fig. 3a-b). To further demonstrate differences in Aβ fluorescence within the two cohorts, we generated high magnification confocal micrographs of six brain regions i.e. striatum (STR), ventral extra-parenchymal CSF space (VCF), lateral extra-parenchymal CSF space (LCF), ependyma (EPM), corpus callosum (CC) and hypothalamus (HTL), as illustrated in Fig. 3a-b. Aβ fluorescence was observed in deeper anatomical regions of the WT mouse brain. On the other hand, Aβ fluorescence in corresponding regions of the *Aqp4* KO mouse brain was dramatically less (Fig. 3c-d). These observations indicate visible sequestration of Aβ peptide close to the striatal site of injection within the *Aqp4* KO mice.

**Fig. 3.**
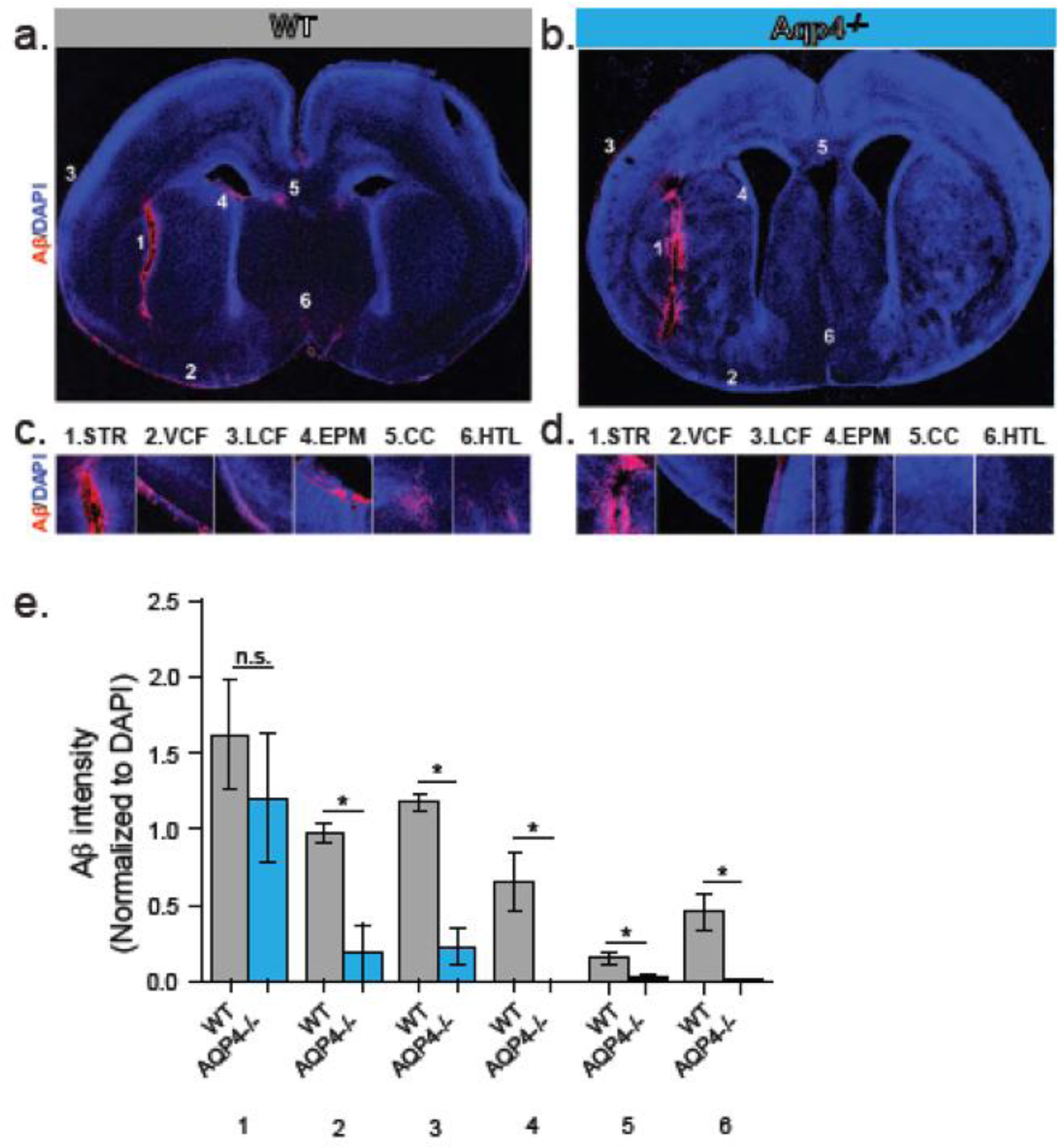
UNC: Comparison of the dispersion of fluorescent Aß after intrastriatal administration in WT and *Aqp4* KO mice. Postnatal day 0 (P0) mice were injected with a fixed concentration (1 μg per animal) of hilyte-555 labeled amyloid-beta (Aß) peptide (AS-60480-01 ; Anaspec, CA) into the left striatum. Three hours post injection, mice were killed by anesthetic overdose, and perfused with 1X PBS followed by 4% PFA, and the brains were harvested, post-fixed and vibratome sectioned. (a-b) Representative whole brain coronal micrograph stitches showing an overlay of fluorescent signals from Aβ peptide (magenta) and nuclear DAPI staining (blue) within the WT and *Aqp4* KO mice. Numbers within the whole brain images indicate the positions of higher magnification insets (below). (c-d) High magnification insets show overlay of fluorescent signals at the following mouse brain regions - striatum (STR), ventral CSF (VF), lateral CSF (LF), ependyma (EPM), corpus callosum (CC) and hypothalamus (HTL). Quantitation of pixel intensities from Aβ peptide (magenta) (normalized to DAPI; blue) within aforementioned brain regions was carried out using ImageJ analysis software. Specifically, the mean intensity calculator function was applied across images from different mouse brain regions. (e) Graphical data shown is represented as mean ± standard deviation. P values were calculated by unpaired, 2-tailed Students t-test. ‘n.s.’ indicates not statistically significant; ‘*’ indicates statistically significant (p <0.05). All experiments were conducted in triplicate, representative images are being shown.

To further support our observations, we performed quantification of Aß fluorescent intensity, normalized to nuclear (DAPI) staining using ImageJ analysis software. No significant difference was observed in normalized Aβ fluorescence intensities between transgenic and WT mice in the area surrounding the striatal site of injection (Fig. 3e, region 1). However, similar quantification showed significantly higher normalized Aβ-fluorescence intensities within other brain regions (VCF, LVF, EPM, CC and HTL) of WT mouse brains compared to *Aqp4* KO mice (Fig. 3e, regions 2-6). Overall, our results show reduced parenchymal spread of Aβ peptide, hours after intra-striatal administration in *Aqp4* KO mouse brains.

### URMC: Cerebrospinal fluid entry to brain occurs along the glymphatic pathway and is facilitated by the presence of AQP4 water channels

In order to evaluate the entry pathways of CSF to the brain we infused Alexa 647-conjugated bovine serum albumin (BSA) into the cisterna magna of anesthetized WT and *Aqp4* KO mice. After 30 min, tracer could be found at the base of the brain around the Circle of Willis of both WT and *Aqp4* null mice (Fig. 4a-e). Tracer evidently entered the brain parenchyma through a network of perivascular spaces of the large cerebral arteries on the pial surface. More tracer could be found on the dorsal cortical surface of WT mice compared to *Aqp4* KO mice (Fig. 4b-e). From the pial surface, tracer primarily entered the brain through perivascular spaces surrounding penetrating arteries in the basal ganglia and cortex, and less tracer uptake was seen in the *Aqp4* KO mice (Fig. 4c-g). The presence of normal AQP4 expression decreased the resistance to CSF entry and increased the distance of tracer around penetrating arteries (Fig. 4d-h); AQP4 deletion reduced CSF entry through the glymphatic pathway by ~42.4% (Fig. 4i). This result is similar to previously reported findings, despite earlier use of a different tracer molecule and infusion paradigm (1). To ensure that the mice used for the present experiments were in fact knockouts, we genotyped using qPCR for both the Aqp4 locus (Fig. 4j) and the excision sequence (Fig. 4k). The results indicate that all mice used for these experiments were either homozygous for the *Aqp4* locus (*Aqp4^+/+^*) or were homozygous knockouts (*Aqp^־/־^*) possessing no copies of exon 1-3 of the *Aqp4* gene.

**Fig. 4.**
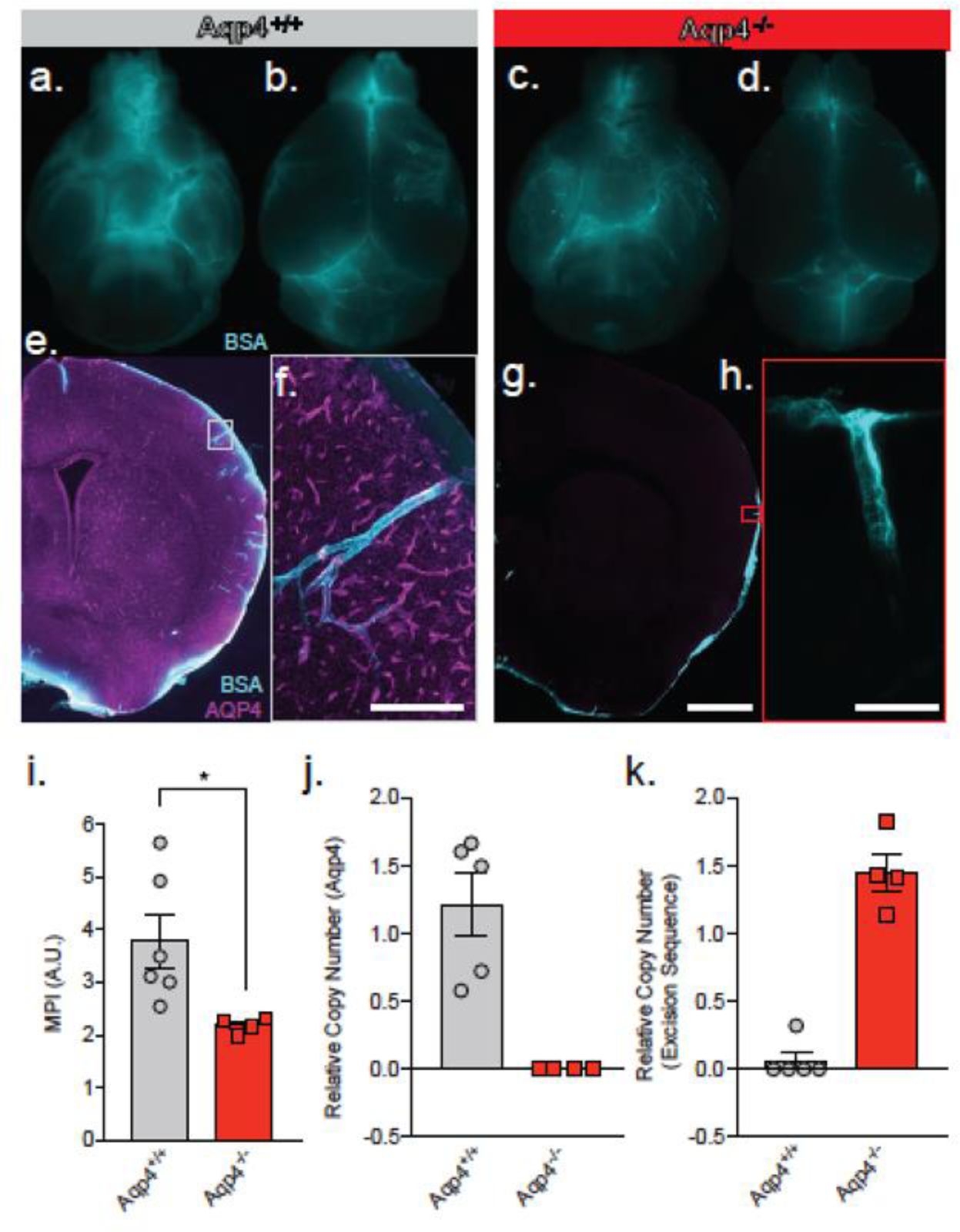
URMC: Glymphatic influx of CSF tracer is facilitated by AQP4. Representative ventral (a) and dorsal (b) whole brain images of WT control mice (*Aqp4^+/+^*) 30 min after intracisternal delivery of a fluorescent tracer, BSA-647. Representative ventral (c) and dorsal (d) whole brain images of *Aqp4* KO (*Aqp^־/-^*) mice. (e) Coronal section from an *Aqp4^+/+^* mouse stained and imaged for AQP4 (magenta) and BSA-647 tracer (cyan). (f) Inset from e (grey box), acquired using high-power confocal laser scanning microscopy showing perivascular entry of the CSF tracer. Scale bar: 150 μm. (g) Coronal section from an *Aqp^־/-^* mouse stained and imaged for AQP4 and intracisternal tracer. Scale bar: 1 mm (h) Inset from g (red box), images acquired using confocal microscopy showing reduced perivascular entry of the CSF tracer and absent AQP4 labeling. Scale bar: 50 μm. (i) Quantification of coronal sections using a non-thresholded approach (mean ± SEM). Data shown as mean pixel intensity (MPI) in arbitrary units (A.U.). *\*P<0.05;* two-tailed unpaired t-test, n=4-6/group. Relative copy numbers (RCN) of the Aqp4 gene locus (j) and the excision sequence (k) showing successful deletion of Aqp4 exon 1-3. RCN was quantified by qPCR for each mouse from experiment in (i).

### OHSU: Reduced glymphatic CSF influx in Sntal KO mice with loss of polarized expression of AQP4 in vascular endfeet of astrocytes

To assess the role of perivascular astrocytic localization of AQP4 in mediating CSF flux into the brain parenchyma, we utilized the *Snta1* KO mouse line. These mice lack expression of the adapter protein α-syntrophin, which links AQP4 to the dystrophin associated complex and is critical for maintenance of the perivascular localization of AQP4 (2). Immunolabeling of AQP4 illustrates the loss of AQP4 perivascular localization in the *Snta1* KO mice (Fig. 5a-b). Higher magnification imaging reveals that, while perivascular localization is lost, widespread AQP4 expression is still detectable by immunofluorescence, consistent with a previous characterization of the mouse line (Fig. 5a-b insets (*48*)). We next sought to determine if parenchymal CSF influx kinetics are altered when perivascular AQP4 localization is lost. Here we used DCE-MRI to measure the influx of the contrast agent gaditeridol (Gad) into brain parenchyma after intracisternal injection. We performed serial T1-weighted imaging at 10 minute intervals following administration of the contrast agent (Fig. 5c-h). At 30 minutes after the start of the infusion, elevated levels of Gad were detected in both WT mice and *Snta1* KO mice, particularly along the ventral surface of the brain (Fig. 5e-f), but by 60 minutes Gad signal within the parenchyma of *Snta1* KO mice was significantly reduced compared to wild type mice (Fig. 5g,h,n). Regional assessment revealed decreased signal in *Snta1* KO mice across brain regions, which was most pronounced in cortex and hippocampus compared to subcortical brain regions and within the ventricles (Fig. 5j-m). Taken together, these data illustrate a role for α-syntrophin in facilitating CSF-ISF exchange and, by extension, support the reported role of AQP4 in this process. Furthermore, these results suggest that the perivascular localization of AQP4 contributes to the kinetics of CSF influx into the brain parenchyma.

**Fig. 5.**
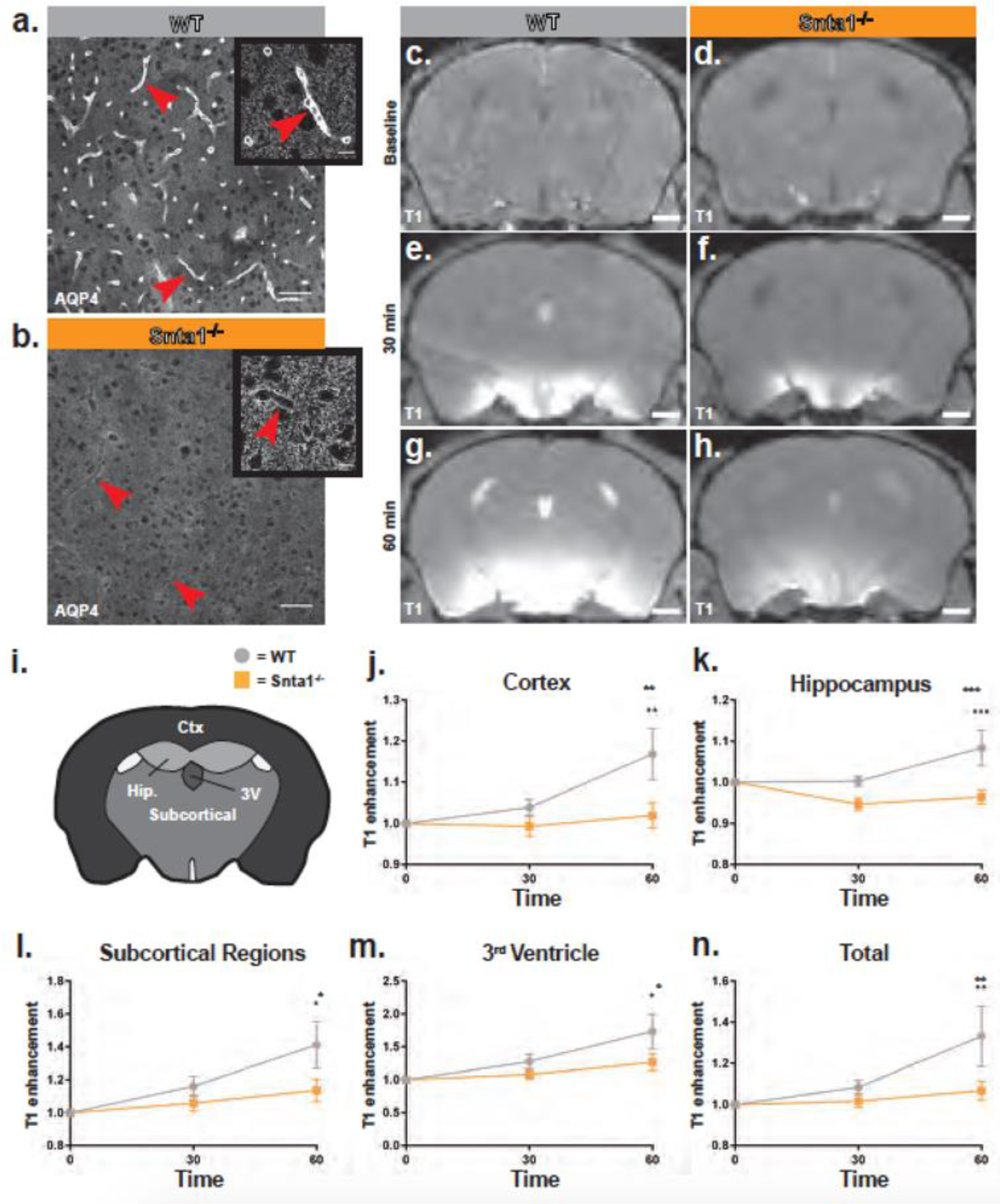
OHSU: Deletion of the adapter protein α-syntrophin impairs AQP4 perivascular localization, and CSF influx into the brain parenchyma. Dynamic contrast enhanced magnetic resonance imaging (DCE-MRI) was used to characterize the effect of α-syntrophin deletion on CSF influx into the brain. Representative images of AQP4 perivascular localization in wild type mice (a), and the loss of perivascular localization of AQP4 seen in the *Snta1* KO mice (b). Scale bar: 50 μm, inset scale bar: 10 μm. (c-h) Coronal slice of T_1_-weighted images acquired by DCE-MRI demonstrate the reduced influx of gaditeridol contrast agent into the parenchyma in *Snta1* KO mice relative to wild type mice at 30 and 60 minutes. Scale bar: 1 mm. (i-n) Quantification of T_1_ weighted signal in various brain sub regions relative to baseline at each time point. (mean ± SEM, 2-way ANOVA). WT n = 5, *Snta1* KO n =7. Ctx = cortex (*p* = 0.0035) Hip = hippocampus (*p* = 0.0003) Subcortical = subcortical regions (*p* = 0.019) 3V = 3^rd^ Ventricle (*p* = 0.028) Total (*p* = 0.0085).

## Acknowledgement

This study was funded by NIH (NS100366, NS078394, NS089709, NS061800, AG048769, AG054456, AG054093, NS099371 and HL089221), NNSF81671070, the Knut and Alice Wallenberg Foundation, and Western Norway Regional Health Authority (Helse Vest). We thank Dan Xue for expert graphic illustrations and Paul Cumming for comments on the manuscript.

